# Auditory evoked potential (P300) in cochlear implant users: a scoping review

**DOI:** 10.1101/2020.05.13.094235

**Authors:** Maria Stella Arantes do Amaral, Nelma Ellen Zamberlan-Amorin, Karina Dal Sasso Mendes, Sarah Carolina Bernal, Eduardo Tanaka Massuda, Miguel Ângelo Hyppolito, Ana Cláudia Mirândola Barbosa Reis

## Abstract

**Introduction:** P300 auditory evoked potential is evoked by a long latency auditory stimulus, which provides information on neural mechanisms underlying central auditory processing, considered an objective and non-invasive technique to study the auditory central nervous system.

**Objective:** To identify and gather scientific evidence regarding the P3 component in adult cochlear implant users.

**Methods:** Scoping review of scientific literature, in the search of original articles in Portuguese, Spanish and English, published between 1991 and May 2018, in the following database websites: PubMed / Medline, Embase, LILACS and Web of Science.

**Results:** A total of 87 articles were identified and exported to the search software Rayyan for study selection - 58 were from Embase, 26 from PubMed and 3 from Web of Science. There were no articles found on LILACS. From those 87 articles, 16 were excluded for being duplicated. Then 71 articles were selected for title, authors, yeas and abstract scanning, from which 50 articles were excluded. From the 21 final articles for full reading, one was excluded for not performing P300, leaving us with 20 selected articles.

**Conclusion:** This review has contributed with evidence that indicates how important it is to include speech stimulation when measuring P300. Regardless of the stimulus being used for P300 elicitation, a pattern of results can be seen a higher latency and a lower amplitude in CI users.

## Introduction

The P300 auditory evoked potential, also known as cognitive potential or event-related potential, is a long-latency cortical potential, evoked by auditory stimulation. This potential is obtained by the registration and mediation of responses to sensorial stimulation captured from a distance, on the skull surface and is obtained through the identification of a rare auditory stimulus presented among other frequent stimuli, known at the oddball paradigm [1,2]. In normal-hearing adult subjects, this potential appears at around 300ms after stimulus presentation, with positive voltage and amplitude between 5 and 20 μvolt [3-6].

Event-related potentials, evoked by auditory stimuli, provide information on neural mechanisms underlying auditory processing. They are considered an objective and non-invasive technique to further study the auditory central nervous system [7-9].

P300 can be achieved in subjects with hearing loss as long as the subject is able to identify the rare stimulus among the frequent ones, and it can be used to monitor hearing impaired patients who are going through rehabilitation, since studies show decreased latency of the P3 component after rehabilitation therapy, highlighting the cognitive improvement of these subjects [10,11].

Studies show a direct association between hearing loss and cognitive capacity changes, indicating that moderate to severe sensorioneural hearing loss can lead to extended latency of N1, N2 and P3 components [12]; or to the impact of hearing loss in the degree of cognitive reduction, as described by other authors [13] or even the influence of hearing loss in cognitive abilities related to word discrimination, comparing the use of P300 in subjects with and without hearing loss [14].

Electrophysiological assessment of the auditory system has been considered an object of study, especially in subjects with hearing loss, who have been deprived of hearing for a period of time and restored their hearing abilities through electric stimulation with a CI. There is a link between auditory deprivation and cognitive function loss, as well as the need to monitor cortical response in face of the new auditory input, as time goes by [15].

Kaga et al. showed us in 1991 that it was possible to achieve P300 potential in a cochlear implant user [16]. The authors performed weekly tests on an adult user of a single-channel CI from the brand House, showing that the P3 component became more robust as the months went by, which indicates this testing could be a useful tool for hearing restoration monitoring, as well as for elucidation of changes in the hearing perception of CI users.

Other authors published studies with different oddball paradigms: tone burst in different frequencies [17-28]; speech stimuli with different contrasting sounds [29-33] and even one study with music paradigm in order to evaluate cortical function in these individuals [34]. On the other hand, authors Makdoum et al, 1998 and Groenen et al, 2001 published studies that used pure tone and speech stimuli [35,36].

Considering P300’s clinical applicability as a possible tool for neuronal plasticity monitoring during CI hearing rehabilitation, our study raised the following question: “What is the scientific evidence in regards to P3 measurements in CI users?”, according to the PCC (Population, Concept and Context) acronym for scoping review [37]. We previously defined the acronym as P: adult subjects with post-lingual hearing loss, C: Cochlear implant surgery and C: P300 examination. To answer such question, our objective was to analyze P3’s latency and amplitude values in CI users (all adults, with post-lingual hearing loss).

It is worth noting that this study’s relevancy is that it offers perspectives that could help us understand the relationships between these measurements and the possibilities of monitoring, decision-making and planning the intervention process, as well as patient and family orientation.

## Materials and methods

### Ethical Aspects

Ethical aspects were thoroughly followed with scientific rigor, appropriate citations and rigorous data treatment and presentation.

### Methodological framework

This study used the methodological approach by *The Joanna Brigs Institute for Scoping Reviews (JBI)* [37].

### Type of study

This is a scoping review, a systematic review which aims to map relevant scientific production in a certain field, in this case, the medical field.

Considering the leading question related to current evidence in the literature on amplitude and latency measurements for the P3 component with different stimuli, speech and pure tone (tone burst) and in regards to clinical applicability for CI users, we then proceeded to search for controlled and not-controlled terms identified in the Medical Subject Headings (MeSH), the National Library of Medicine (NLM) and the Health Science Descriptors (DeCS).

The following structure was used to develop a search strategy, PCC: Population – post-lingual adults, Concept – CI surgery, Context – P300 results comparison (Table 1) [39]. The search for original articles was conducted in the following databases: PubMed/Medline, Embase, Lilacs, Web of Science, implemented according to each database criteria and manuals.

**Table 1.** Research methodology with the use of the PCC strategy.

| Population | Concept | Context |
| --- | --- | --- |
| Post-lingual adults | Cochlear Implant | P300<br>Behavior |
| Post-lingual |  | P300 |

As research strategy descriptors, the following words were used: Adult, Adults, post-linguals, post-lingual, Cochlear Implants, Implants, Cochlear, Cochlear Implant, Implant, Cochlear Prosthesis, Cochlear Prostheses, Prostheses, Prosthesis, Auditory Prosthesis, Auditory Prostheses, Prostheses, Auditory, Prosthesis, Auditory, Event-Related Potentials, P300, Event Related Potentials, P300, Event-Related Potential, P300, P300 Event-Related Potential, P300 Component, P300 Components, Event-Related Potentials, P3, Event Related Potentials, P3 Event-Related Potentials, Event-Related Potential, Evoked Potentials, P300 Component, P300 Event-Related Potentials, P300 Event Related Potentials, P3b Event-Related Potentials, Event-Related Potential, P3b, Event-Related Potentials, P3b Event Related Potentials, P3a Event-Related Potentials, P3a, P3a Event Related Potentials, P300, Potenciales Relacionados con Evento P300, Potencial Evocado P300, Componente P300 de Potencial Evocado; combinados com operadadores booleanos (AND e OR), as seen on Table 2.

**Table 2.** Cross-check data from PubMed Medline, Embase, LILACS, Web of Science.

| Data Base | Strategy |  |  |
| --- | --- | --- | --- |
|  | Population | Concept | Context |
| **Pubmed,**<br>**Web of**<br>**Science,**<br>**Embase** | "Adult"[Mesh]<br>OR "Adults"<br>OR "post-linguals" OR<br>"post-lingual" | "Cochlear Implants"[Mesh] OR<br>"Implants, Cochlear" OR "Cochlear<br>Implant" OR "Implant, Cochlear"<br>OR "Cochlear Prosthesis" OR<br>"Cochlear Prostheses" OR<br>"Prostheses, Cochlear" OR<br>"Prosthesis, Cochlear" OR<br>"Auditory Prosthesis" OR<br>"Auditory Prostheses" OR<br>"Prostheses, Auditory" OR<br>"Prosthesis, Auditory" | "Event-Related Potentials, P300"[Mesh] OR "Event Related Potentials, P300"<br>OR "Event-Related Potential, P300" OR "P300 Event-Related Potential" OR<br>"P300 Component" OR "P300 Components" OR "Event-Related Potentials,<br>P3" OR "Event Related Potentials, P3" OR "P3 Event-Related Potentials" OR<br>"Event-Related Potential, P3" OR "P3 Event Related Potentials" OR "P3<br>Event-Related Potential" OR "Evoked Potentials, P300 Component" OR<br>"P300 Event-Related Potentials" OR "P300 Event Related Potentials" OR<br>"P3b Event-Related Potentials" OR "Event-Related Potential, P3b" OR<br>"Event-Related Potentials, P3b" OR "P3b Event Related Potentials" OR "P3b<br>Event-Related Potential" OR "P3a Event-Related Potentials" OR "Event-Related Potential, P3a" OR "Event-Related Potentials, P3a" OR "P3a Event<br>Related Potentials" OR "P3a Event-Related Potential" |
| **LILACS** | Cochlear<br>Implantation<br>OR<br>Implantación<br>Coclear OR<br>Implante<br>Coclear OR<br>Implantação<br>Coclear OR<br>Implante de<br>Prótese<br>Coclear | Cochlear Implantation OR<br>Implantación Coclear OR Implante<br>Coclear OR Implantação Coclear<br>OR Implante de Prótese Coclear | Event-Related Potentials, P300 OR Potenciales Relacionados con Evento<br>P300 OR Potencial Evocado P300 OR Componente P300 de Potencial<br>Evocado |

The research strategy was implemented in a standardized way for each database, with adaptations when needed. Files were exported to the reference manager EndNote, version X5, for removal of duplicates. Then, a new file was created, which was then exported to the Rayyan software, a specific tool for study selection in review methods [40].

Selection criteria consisted in the search for studies in Portuguese, Spanish and English, published between January 1991 and May 2018. We included in this search articles that studied adult subjects, with pos-lingual hearing loss, who were submitted to cochlear implantation surgery and were tested with P300. We excluded the following: case reports, reviews, articles in press, letters to the editor and studies in other languages that are not dominant to the researchers.

The following flow chart represents the path of identification, selection, inclusion of selected primary studies, following the consulted electronic database (fig 1), regardless of evidence level.

**Fig 1.**
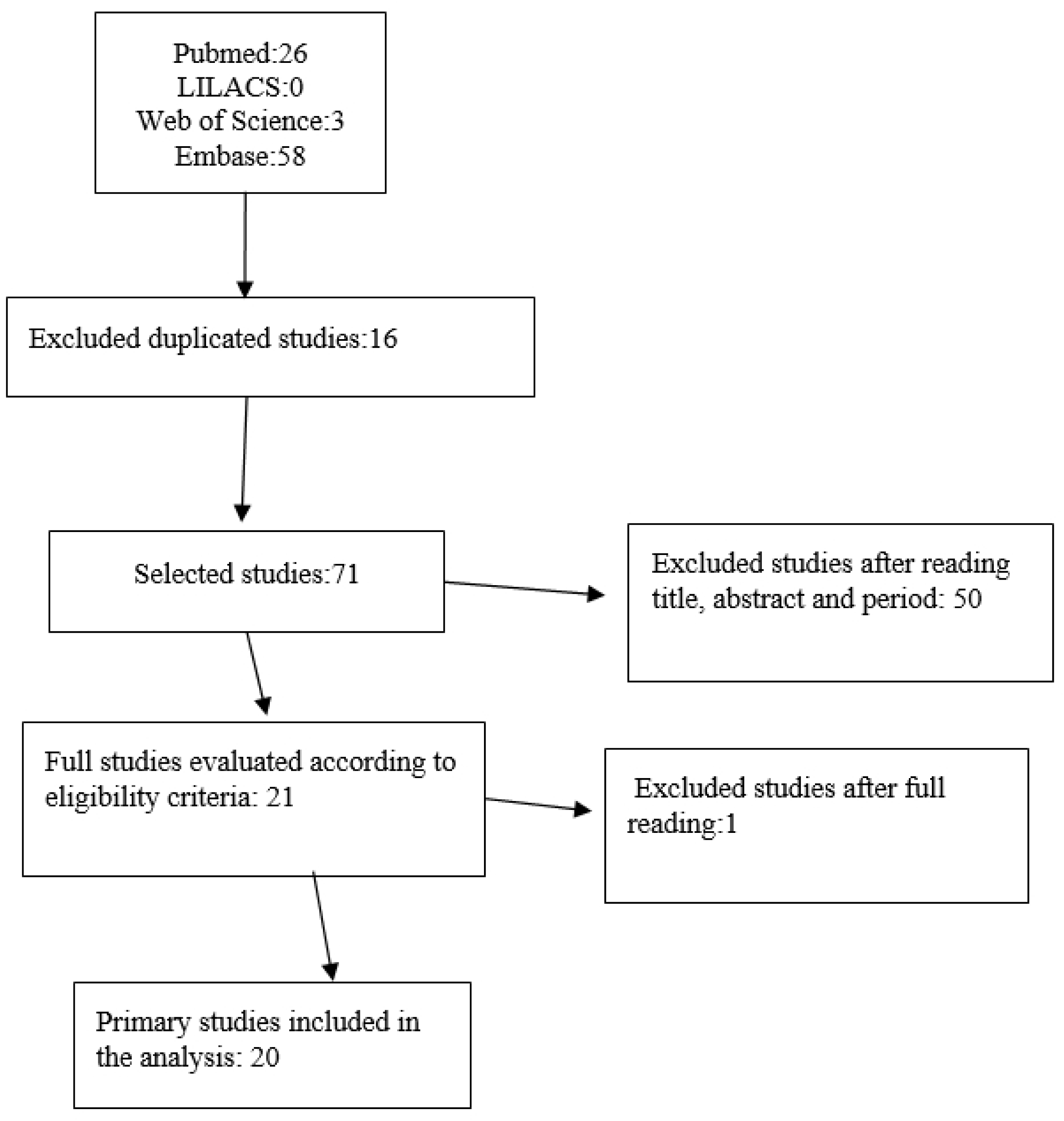
Flow-chart of identification, selection and inclusion of studies from the integrative review.

The process of study selection was performed by two reviewers with an independent method, following previously established inclusion and exclusion criteria. On the first phase, they performed reading of title and abstract, following the established inclusion and exclusion criteria, more articles were excluded. A meeting for consensus regarding divergencies related to study inclusion took place. Then on the second phase, texts were fully read and excluded according to the same criteria. A percentage of interrater reliability between the reviewers (90%) was used and, in case of disagreement, a third reviewer was invited.

For data extraction, a standardized script was used, with characterization of each study (author, year, methodological aspects and main results), and data related to the type of stimulus used and specified according to P300 measurements.

For results presentation, descriptive data analysis was conducted, including tables with the studies’ synthesis, searching for answers regarding latency and amplitude measurements found in each article, as well as the variables and parameters identified between them. We used the Preferred Reporting Items for Systematic Reviews and Meta-Analyses (PRISMA) to report data [38].

## Results

A total of 87 articles on the Rayyan software - 58 from the Embase database, 26 from PubMed and 3 from the Web of Science platform were selected for this study. No articles were located in the LILACS database. From those 87 articles, 16 were excluded for duplicity. Then 71 articles were selected for title, authors, year and abstract reading, from which 50 were excluded. From the 21 remaining articles, one was excluded for not containing P300 testing, leaving us with 20 selected articles.

From those selected 20 original articles, four were published in 2004, there were two articles for each of the following years: 2014, 2009, 2005, 2001 and one article for each of these years: 2018, 2016, 2015, 2012, 2007, 1999, 1998, 1996. (E1 to E20, Table 3) [17-36].

**Table 3.** Synthesis of the primary studies, presented in order of stimulus used, year of publication, author, title and objective.

| STUDY | STIMULUS | YEAR | AUTHORS | TITLE | OBJECTIVE |
| --- | --- | --- | --- | --- | --- |
| E1 | Pure tone | 1996 | Groenen,P.A.P. et al.<sup>17</sup> | The relation between electric auditory brain stem and cognitive responses and speech perception in cochlear implant users | To correlate results for short and long latency potentials with speech perception tests in CI users. |
| E2 | Pure tone | 1997 | Jordan,K. et al<sup>18</sup> | Auditory event-related potentials in post- and prelingually deaf cochlear implant recipients | To observe P300 behavior in CI users 6 months after activation. |
| E3 | Pure tone | 1999 | Okusa, M., et al.<sup>19</sup> | Effects of discrimination difficulty on cognitive event-related brain potentials in patients with cochlear implants | To investigate the effects of discrimination difficulty with 4 stimuli conditions in CI users. |
| E4 | Pure tone | 2001 | Kubo, T., et al.<sup>20</sup> | Significance of auditory evoked responses (EABR and P300) in cochlear implant subjects. | To examine the significance of remaining ganglionic neurons, with auditory evoked potentials and correlate that with speech perception tests in CI users. |
| E5 | Pure tone | 2004 | Iwaki, T., et al<sup>21</sup> | Comparison of speech perception between monaural and binaural hearing in cochlear implant patients | To evaluate the advantages of binaural hearing for unilateral and bilateral CI users, through tests such as P300. |
| E6 | Pure tone | 2004 | Muhler, R., et al<sup>22</sup> | Visualization of stimulation patterns in cochlear implants: application to event-related potentials (P300) in cochlear implant users. | To demonstrate the effect of stimulation pattern variation in P300 in CI users. |
| E7 | Pure tone | 2005 | Kelly, A. S., et al.<sup>23</sup> | Electrophysiological and speech perception measures of auditory processing in experienced adult cochlear implant users | To determine the relationship between evoked potentials and speech perception tests in CI users. |
| E8 | Pure tone | 2007 | Nager, W., et al.<sup>24</sup> | Automatic and attentive processing of sounds in cochlear implant patients - electrophysiological evidence. | To evaluate whether CI users' difficulties to rare stimuli is due to an attention deficit. |
| E9 | Pure tone | 2009 | Sasaki, T., et al.<sup>25</sup> | Assessing binaural/bimodal advantages using auditory event-related potentials in subjects with cochlear implants | To evaluate the advantage of binaural and bimodal hearing for CI users through evoked potentials and speech perception tests. |
| E10 | Pure tone | 2012 | Obuchi, C., et al.<sup>26</sup> | Auditory Evoked Potentials under Active and Passive Hearing Conditions in Adult Cochlear Implant Users. | To investigate the relationship between P300 and MMN using active and passive hearing paradigms with CI users. |
| E11 | Pure tone | 2015 | Finke, M., et al.<sup>27</sup> | Auditory distraction transmitted by a cochlear implant alters allocation of attentional resources | To analyze cortical responses in different stages and correlate them with visual and auditory distractors in CI users. |
| E12 | Pure tone | 2018 | Grasel, S. .et al<sup>28</sup> | P3 Cognitive Potential in Cochlear Implant Users | To assess changes in P300 latency and amplitude in CI users. |
| E13 | Speech | 2005 | Beynon, A. J., et al.<sup>29</sup> | Discrimination of Speech Sound Contrasts Determined with Behavioral Tests and Event-Related Potentials in Cochlear Implant Recipients | To study P300 variation in CI users with different contrasting sounds as stimulus. |
| E14 | Speech | 2009 | Henkin, Y., et al.<sup>30</sup> | Cortical Neural Activity Underlying Speech Perception in Postlingual Adult Cochlear Implant | To examine the relationship between P300 and behavior measurements. |

|  |  |  |  | Recipients |  |
| --- | --- | --- | --- | --- | --- |
| **E15** | Speech | 2014 | Soshi et al.<sup>31</sup> | Event-related potentials for better speech perception in noise by cochlear implant users | To investigate neurophysiological and behavioral aspects for speech perception in noise by CI users. |
| **E16** | Speech | 2014 | Henkin, Y., et al.<sup>32</sup> | Neural correlates of auditory-cognitive processing in older adult cochlear implant recipients." | To compare P300 in elderly CI users and normal-hearing older adults. |
| **E17** | Speech | 2016 | Finke, M., et al.<sup>33</sup> | On the relationship between auditory cognition and speech intelligibility in cochlear implant users: An ERP study | To investigate auditory and cognitive processing for words presented in different conditions and related to cognitive and speech intelligibility abilities in CI users. |
| **E18** | Pure tone e<br>Speech | 1998 | Makhdoum, M. J.<br>A., et al<sup>35</sup> | Can event-related potentials be evoked by extra-cochlear stimulation and used for selection purposes in cochlear implantation? | To investigate whether P300 responses can be elicited by extra-cochlear stimulation using tone and speech stimuli. |
| **E19** | Pure tone e<br>Speech | 2001 | Groenen, P. A. P.,<br>et al <sup>36</sup> | Speech-evoked cortical potentials and speech recognition in cochlear implant users | To correlate P300 results with behavior test results and speech perception. |
| **E20** | Musical | 2004 | Koelsch, S. et al<sup>34</sup> | Music perception in cochlear implant users: an event-related potential study. | To compare music irregularities and physical oddballs between CI users and normal-hearing individuals. |

**Table 4.**
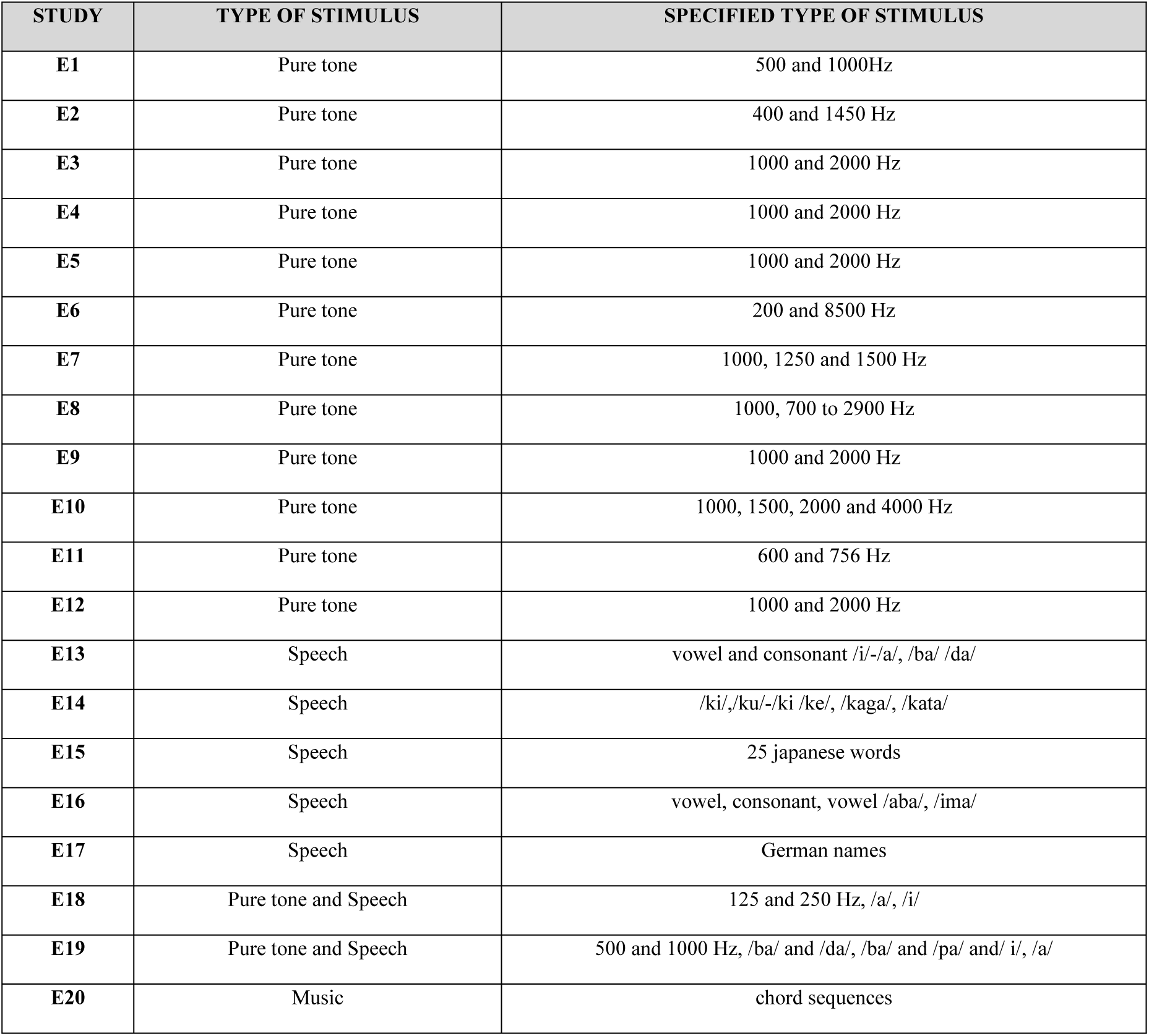
Synthesis of article abstracts, specified by type of stimulus and detailed specifications.

As for the origin of these studies, they were all published in English, 19 of them were published in international journals and only one was published in a Brazilian national journal, revealing the lack of national articles about this subject.

In regards to the parameters used to elicit P300, such as the details on which stimulus was used, these data are further described in chart 4. Twelve articles used pure tone stimuli (E1 to E12) [17-28], five articles used speech stimuli (E13 to E17) [29-33], one study used musical stimulation (E20) [34] and two articles used speech and pure tone stimuli (E18 and E19) [35,36].

Considering that sample size and stimuli can influence P3 registrations, specifically in comparison to latency and amplitude measures, we sought to demonstrate results regarding these variables (Table 5).

**Table 5.** Synthesis of article abstracts, specified by case-by-case subject analysis and obtained results.

| STUDY | SUBJECTS | RESULTS |
| --- | --- | --- |
| E1 | 7 CI users and 11 normal-hearing | P300 latencies were decreased in good CI users, compared to average ones.<br>Average CI users showed increased latency when compared to normal-hearing individuals. |
| E2 | 13 CI users | Latency increased and amplitude decreased as the task's difficulty increased. |
| E3 | 8 CI users | Latency increased and amplitude decreased as the rare stimulus approaches the frequent ones. |
| E4 | 25 CI users e 25 normal-hearing | Higher P300 latency in subjects with lower speech perception scores. |
| E5 | 6 CI users | Higher P300 latency in subjects with lower speech perception scores. |
| E6 | 2 CI users | Amplitude was reduced when the task's difficulty increased. |
| E7 | 12 CI users and 12 normal-hearing | Amplitude was higher and latency lower as time of experience with CI increased. |
| E8 | 7 CI users and 7 normal-hearing | P300 amplitude in the passive condition was lower than in active condition. CI users showed reduced amplitude when compared to control. |
| E9 | 15 CI users, 4 bilateral and 11 bimodal | Latency was lower in binaural subjects compared to monaural subjects. |
| E10 | 3 CI users e 3 normal-hearing | Amplitude levels were lower and latency levels were higher in CI users. |
| E11 | 12 CI users e 12 normal-hearing | P300 latency was higher and amplitude was reduced in the presence of visual and sound distractors. |
| E12 | 26 CI users e 26 normal-hearing | P300 latency was similar to the control group in CI users with a good speech perception performance. |
| E13 | 10 CI users e 10 normal-hearing | P300 amplitudes were reduced and latency were increased for vowel and consonant contrast compared to the control group. |
| E14 | 15 CI users e 12 normal-hearing | P300 latency was similar to the control group in CI users with better speech perception test performance. |
| E15 | 17 CI users e 12 normal-hearing | Higher amplitude levels were correlated to higher speech perception test performance. |
| E16 | 9 CI users e 10 normal-hearing | P3 prolonged and decreased in CI users with age above 60. |
| E17 | 13 CI users e 13 normal-hearing | P300 prolonged in CI users exposed to noise. |
| **E18** | 5 extra cochlear CI and 9 intra | Latency levels were prolonged in extra and intra cochlear CI users, in |
|  | cochlear | comparison to normal-hearing individuals. Results with pure tone were similar and there was correlation between amplitude and speech perception. |
| E19 | 9 CI users e 10 normal-hearing | A relationship was found between P300 amplitude for 500 and 1000Hz and /a/./i/ and speech perception tests. |
| E20 | 12 CI users e 12 normal-hearing | P300 present with music stimulation, with reduced amplitude and increased latency. |

## Discussion

From the selected articles it was possible to identify that the methodologies proposed were not homogeneous. Different criteria are associated to the protocol used and the parameters for P300 testing, according to the objective of each study, once the exam’s parameters should meet the need to investigate. Most of the articles (60%) adopted pure tone as presented stimulus for P300, whereas five articles (25%) used speech, two (10%) used speech and pure tone, and only one used music stimulation (Table 3). We can understand why the option of studying cortical auditory potentials with a speech stimulus is so important, once the goal in this kind of intervention is to give the patient access to speech sounds. Another important factor to consider is the advantage in associating objective tests, such as electrophysiological tests, to behavioral assessment (like speech perception tests), as a novel resource to understand the auditory system and also the limitations of neuronal plasticity and its consequences in speech perception performance. This perception of the need to adjust a pattern in parameters - in this case, the stimuli – for auditory evoked potentials has had an increased importance over the last decade, which can be seen in this study, with publications from 2005 on (Table 3, E13 to 17) [29-33].

Another factor which has been widely discussed in literature and that can be seen in this review is the variability of established parameters for P300 registration [7]. Nonetheless, many studies in this review agree that P300 results contribute with strong evidence for complex interactions between speech intelligibility [31-33], neural processing [30,32], verbal working memory and subjective classifications of hearing effort in cochlear implant users [19,24].

However, although heterogeneity can be observed, we found as a result in 60% of this review (Table 5) studies that compare CI users with normal-hearing individuals, with data that show increased latencies of the P3 component in CI users (E1, E2, E3, E4, E5, E10, E11, E13, E16, E17, E18, E20) and decreased amplitude values of P3 in CI users, in 35% of the studies (E6, E8, E10, E11, E13, E16, E20). These findings make us think of a few hypotheses, such as the influence of de CI’s external component, speech processor and programming options on latency increase. The second hypothesis (which does not exclude the first one) are the intrinsic aspects of each individual that could interfere in these results, for example the listening effort of the hearing-impaired subject and cognitive aspects inherent to hearing abilities, especially related to attention and memory.

Attention is an important aspect for this test since the P3 component is registered when the patient notices the rare stimulus. Memory seems to be related to the solicited task during the test (to mentally count), which in most publications isn’t explicit, since only 4 studies (20%) out of 20 explained that subjects were instructed to count the rare stimulus (E1, E12, E19 and E20) [28, 34, 36].

In the E7 [23] study, we could notice that a higher time of hearing experience using the CI was associated with a lower latency of P300 components. These cumulative results suggest that pathway maturation is possible with an increased hearing experience and that a maximum level of central auditory pathway maturation could occur before a second CI, which leads us to reflect on the moment of surgery and the fact that it could influence results with the second CI, especially in children with sequential bilateral CI [41].

An important aspect found in this review were the results of latency measurements of the P3 component being similar between CI users and normal hearing individuals, even after a long time of auditory deprivation (E12, E14, E15). Amplitude measurements were also similar between CI users and normal hearing subjects (E18, E19), which suggests these results are related to better results on speech tests. Therefore, it all leads us to think that, in adults with hearing impairment, auditory pathways can remain functional for a long period of time and the central auditory system could be preserved, even when conventional hearing aids don’t provide the ideal auditory stimulation.

As we analyze the influence of the kind of stimulus used to elicit P300, the absolute latency of P3 was considered prolonged in the cortical potential exam on six out of 11 studies that used pure tone (E2^18^ E3^19,^ E4^20,^ E5 E10, E17). When pure tone and speech were both used, study E18 [35] found prolonged latency of the P3 component also.

Age was an aspect highlighted by study E16 as a possible reason for a rise in absolute latency (Henkin, Y., et al.,2014) [32].

Absolute latency may be associated with time of CI experience and some of the studies that used pure tone showed that, with more time of experience, the absolute latency showed reduced levels (E1 E7 and E9).

Amplitude was also a parameter of interest in this integrative review. There were no differences between the amplitude of P3 for monaural and binaural conditions (E5). However, it was one of the parameters which showed correlation between pure tone and speech stimuli and the speech perception test results (E18 and E19).

Hearing loss etiology should also be considered when higher P3 component latencies are found and when there are poor speech perception results, as identified in E12^28^, with meningitis patients and, therefore, authors suggest neuropsychological evaluation in the test battery for CI indication, as a measurement to prevent poor prognosis in CI results.

The effect of music stimulation stood out among other stimuli used to elicit P300, although only one study in this review used this method (E20) [34], showing reduced amplitude and increased latency with this condition. Authors point out that the presence of music-related effects in CI users show that they still have a representation of system regularities, even after a long period of auditory deprivation, in spite of the auditory input provided by a CI.

## Conclusions

This review has contributed with evidence that show how important it is to include speech stimulation in P300.

Regardless of which stimulus was used to elicit P300, we can notice a pattern in latency increase and amplitude decrease in CI users.

Experienced time of CI use and the speech tests’ results seem to be related to the latency and amplitude measurement results of the P3 component.

## Acknowledgements

The authors would like to thank Maria Cecilia Onofre for text corrections and review.

## Supporting information

**S1 Fig. Figure 1. Flow-chart of identification, selection and inclusion of studies from the integrative review**.

**S1 Table 1. Research methodology with the use of the PICO strategy**.

**S2 Table 2. Cross-check data from PubMed Medline, Embase, LILACS, Web of Science**.

**S3 Table 3. Synthesis of the primary studies, presented in order of stimulus used, year of publication, author, title and objective**.

**S4 Table 4. Synthesis of article abstracts, specified by type of stimulus and detailed specifications**.

**S5 Table 5. Synthesis of article abstracts, specified by case-by-case subject analysis and obtained results**.

